# An Integrated Systems Biology Approach Identifies the Proteasome as a Critical Host Machinery for ZIKV and DENV Replication

**DOI:** 10.1101/2020.03.04.976548

**Authors:** Guang Song, Emily M. Lee, Jianbo Pan, Miao Xu, Hee-Sool Rho, Yichen Cheng, Nadia Whitt, Shu Yang, Jennifer Kouznetsova, Carleen Klumpp-Thomas, Samuel G. Michael, Cedric Moore, Ki-Jun Yoon, Kimberly M. Christian, Anton Simeonov, Wenwei Huang, Menghang Xia, Ruili Huang, Madhu Lal-Nag, Hengli Tang, Wei Zheng, Jiang Qian, Hongjun Song, Guo-li Ming, Heng Zhu

**Affiliations:** Department of Pharmacology & Molecular Sciences; Johns Hopkins School of Medicine, Baltimore, MD 21205, USA; Department of Biological Science, Florida State University, Tallahassee, FL, 32306, USA; Department of Ophthalmology; Johns Hopkins School of Medicine, Baltimore, MD 21205, USA; National Center for Advancing Translational Sciences, National Institutes of Health, Bethesda, MD 20892, USA; Sir Run Run Shaw Hospital, Zhejiang University School of Medicine, Hangzhou, 310016, China; Institute for Cell Engineering, Johns Hopkins University School of Medicine, Baltimore, MD 21205, USA; Department of Neuroscience and Mahoney Institute for Neurosciences, Perelman School for Medicine, University of Pennsylvania, Philadelphia, PA 19104, USA; Department of Cell and Developmental Biology, Perelman School for Medicine, University of Pennsylvania, Philadelphia, PA 19104, USA; Institute for Regenerative Medicine, Perelman School for Medicine, University of Pennsylvania, Philadelphia, PA 19104, USA; The Epigenetics Institute, Perelman School for Medicine, University of Pennsylvania, Philadelphia, PA 19104, USA

**Keywords:** Protein-protein interaction, RNAi screening, Chemical genetics screening

## Abstract

The Zika (ZIKV) and dengue (DENV) flaviviruses exhibit similar replicative processes but distinct clinical outcomes. A systematic understanding of virus-host protein-protein interaction networks can reveal cellular pathways critical to viral replication and disease pathogenesis. Here we employed three independent systems biology approaches toward this goal. First, protein array analysis of direct interactions between individual ZIKV/DENV viral proteins and 20,240 human proteins revealed multiple conserved cellular pathways and protein complexes, including proteasome complexes. Second, an RNAi screen of 10,415 druggable genes to identify host proteins required for ZIKV infection uncovered proteasome proteins. Third, a high-throughput screening of 6,016 bioactive compounds for ZIKV inhibitors yielded 134 effective compounds, including six proteasome inhibitors that suppress both ZIKV and DENV replication. Integrative analyses of these orthogonal datasets pinpoints proteasome as critical host machinery for ZIKV/DENV replication. Our study provides multi-omics datasets for further studies of flavivirus-host interactions, disease pathogenesis, and new drug targets.

## Introduction

Dengue virus (DENV) and Zika virus (ZIKV) are two closely related pathogens of the Flaviviridae family [1]. Although dengue disease has been recognized in the Americas since the 1600’s, DENV was only isolated in 1943 and is still one of the most widespread global mosquito-borne viruses, contributing to symptoms in 96 million people and over 20,000 deaths each year [2, 3]. ZIKV was first discovered as a mild, obscure human pathogen in 1947, but has emerged as a major public health concern in the past few years due to its role as an etiological agent in several neurological pathologies, including congenital microcephaly and Guillain-Barre syndrome [4].

The genome of both DENV and ZIKV are composed of a single positive-strand RNA, which is directly translated into a polyprotein and subsequently processed to generate components necessary for viral replication and assembly [1]. Because of the limited number of proteins encoded by viral genomes, viruses are obligatory intracellular pathogens and completely dependent on their hosts for survival and reproduction, which is mediated by direct interactions between the virus and host cellular components [5–7]. A better understanding of virus-host interactions can reveal critical cellular pathways that are necessary for viral replication and for pathogenesis, which in turn could be used to identify effective treatment regimens targeting host proteins [5, 7–9]. Advancements in high-throughput technologies over the last decade have made it possible to systematically analyze the protein-protein interactome between a virus and its host [10–14]. Previous studies have identified several new host pathways that are essential to life cycles of several pathogens, including Kaposi’s sarcoma-associated herpesvirus [15–18], influenza virus [19], HIV [20], and Epstein-Barr virus [21].

Most antiviral drugs are classified as direct-acting antivirals (DAAs). DAAs directly target specific viral proteins critical for infection. While there are many successful DAAs currently in use for viral infections (e.g. hepatitis C virus), it is well-known that many RNA viruses rapidly develop drug resistance due to the selective stress imparted by targeting essential viral proteins and the high mutation rate in their RNA-based genomes [22, 23]. On the other hand, a drug targeting critical host proteins would provide a higher genetic barrier for a virus to develop drug resistance [6].

Genetic similarities between DENV and ZIKV, together with recent findings about the host cell dependency factors they shared, suggest that these two related flaviviruses likely utilize a similar replicative strategy in the host [24, 25]. Consequently, characterization of conserved flavivirus-human protein-protein interactions (PPIs) can reveal critical cellular pathways that are essential for flavivirus infection [5, 25, 26]. On the other hand, differences in PPIs between ZIKV and DENV may provide insight into how these two viruses lead to different pathological outcomes, for example, microcephaly induced by ZIKV [27]. Here, we comprehensively surveyed the human proteome with individual ZIKV and DENV proteins to identify virus-host PPI networks. Bioinformatics analyses revealed multiple cellular pathways and protein complexes, including the proteasome complex. In parallel, a RNAi screen targeting druggable genes and a high-throughput chemical genetics approach also revealed overlapping cellular pathways and protein complexes. Through integrative analysis of these three omics datasets, we identified several conserved cellular machineries important for ZIKV and DENV infection, including the proteasome pathway. Cell-based assays confirmed that proteasome inhibitors effectively suppressed both ZIKV and DENV replication. Together, our study not only provides a valuable multi-omics dataset resource for the field, but also suggests a new strategy for understanding molecular mechanisms of virushost interactions and pathogenesis, and for identifying cellular host-based targets to develop antiviral therapeutics.

## Results

### ZIKV and DENV recombinant proteins

The flavivirus genome encodes a total of ten proteins, including three structural proteins and seven non-structural proteins. Three structural proteins are capsid protein (C), the pre-membrane protein (prM), which is subsequently cleaved upon viral maturation into a Pr peptide and a mature membrane protein (M), and the envelope protein (E), which mediates fusion between viral and cellular membranes. Seven non-structural proteins are non-structural protein 1 (NS1), which is required for formation of the replication complex and recruitment of other non-structural proteins to the ER-derived membrane structures; non-structural protein 2A (NS2A), involving into virion assembly and antagonizes the host alpha/beta interferon antiviral response; Serine protease NS2B (NS2B); serine protease NS3 (NS3); non-structural protein 4A (NS4A), regulating the ATPase activity of the NS3 activity; non-structural protein 4B (NS4B), inducing the formation of ER-derived membrane vesicles; and RNA-directed RNA polymerase NS5 (NS5), as well as the short peptide 2k [28]. To construct ZIKV- and DENV-host PPI networks, we cloned the genes encoded by ZIKV MR766 strain (African strain) and DENV serotype 1 (Figure S1A). We confirmed cloning fidelity by Sanger sequencing (Figure S1B-D and Table S1). Using a previously reported protocol [21], the viral proteins were purified from yeast as N-terminal tagged GST fusion proteins and fluorescently labeled individually (Figure S1E). The quality and quantity of these labeled proteins was evaluated on sodium dodecyl sulfate polyacrylamide gel electrophoresis (Figure S1E).

Considering the varied post-translational modifications (i.e. glycosylation) catalyzed by yeast cells and importance of correct disulfide bond formation involving into protein’s function and binding activity, we decided to focus on the six homologous non-structural proteins (i.e., NS2A, NS2B, NS3, NS4A, NS4B, NS5) and two variants with the signal peptide 2K (i.e., NS4A+2K and 2K+NS4B) encoded by ZIKV (MR766 strain) and DENV-I to construct host-viral PPI networks.

### Construction of ZIKV- and DENV-human PPIs with HuProt™ array

The Human Proteome Microarray v3.0 (HuProt™ array), comprised of 20,240 immobilized human proteins from >15,000 full-length genes, was used to identify human-viral PPI networks [29]. Each viral protein was fluorescently labeled and individually probed to the HuProt array. Fluorescent signals indicating viral protein bound to immobilized human protein were acquired, normalized, and quantified [30]. We used a very stringent cut off (Z-score ≥ 15) to identify positive hits for each viral protein (**Figure 1**A). Examination of assays performed in duplicate showed high reproducibility as measured by Pearson correlation coefficients. An example of binding signals obtained with DENV NS5 is shown in Figure 1B.

**Figure 1.**
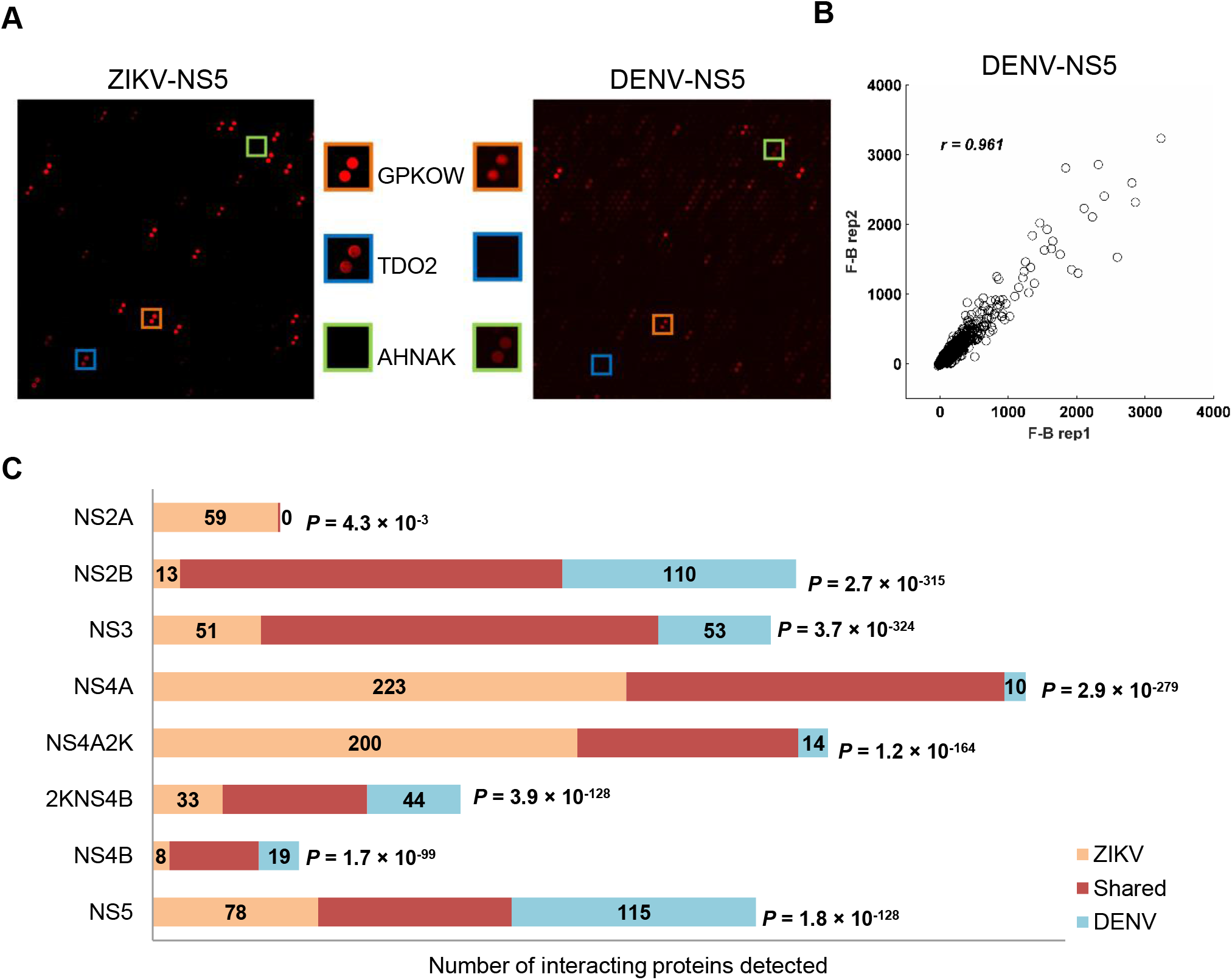
Identification of ZIKV- and DENV-host protein-protein interactomes by protein array. **A.** Sample images of HuProt arrays showing human proteins bound by individual viral proteins. Each human protein was printed in duplicate. The orange, blue and green boxes represent shared (red), ZIKV-(blue), and DENV-(green) specific interactions. **B**. Duplicate experiments performed for each virus protein probe showed high reproducibility. Pearson correlation coefficient analysis showed signals of duplicate experiments based on DENV-NS5 have a high linear relationship, ***r*** =0.961. **C.** Summary of numbers of unique and conserved virus-host interactions between each ZIKV and DENV homologous pair.

We identified a total of 1,708 ZIKV-host PPIs and 1,408 DENV-host PPIs, involving 581 human proteins (Table S2). The majority of host proteins were found to interact with specific individual viral proteins. For example, 152 human proteins only interacted with a single viral protein, whereas 75 human proteins bound to two viral proteins. We found 24 human proteins that interacted with all viral proteins tested (Figure S2A), possibly a consequence of the common N-terminal GST tag. These 24 human proteins were removed from further analysis.

Actually, we recently used the NS2A PPI dataset to investigate how ZIKV-NS2A causes microcephaly-like phenotypes in the embryonic mouse cortex and human forebrain organoid models [27]. Using a co-immunoprecipitation assay, we confirmed interactions between ZIKV-NS2A and several endogenous PPI targets (ARPC3, SMAD7, NUMBL) in neural stem cells [27]. We also evaluated the ability of our approach to recover human proteins known to be targeted by viruses. We acquired a total of 754 human targets from VirusMint [31] and Virhostome [32]. Of the 581 host proteins identified in our PPI analysis, 54 overlapped with the 754 VirusMint or Virhostome targets (hypergeometric *p*-value = 1.9e-5; Fig. 2B). Also, this around 10% identified hits might be caused by different technologies and varies of virus have their specific binding proteins to maintain their replication. Furthermore, we noted that a near published paper identified 701 vs. 688 human binding proteins by IP-MS and BioID respectively (Fig.2C), both of which were based on MS. Of them, 48 overlapped with our data (hypergeometric *p*-value = 0.004).

**Figure 2.**
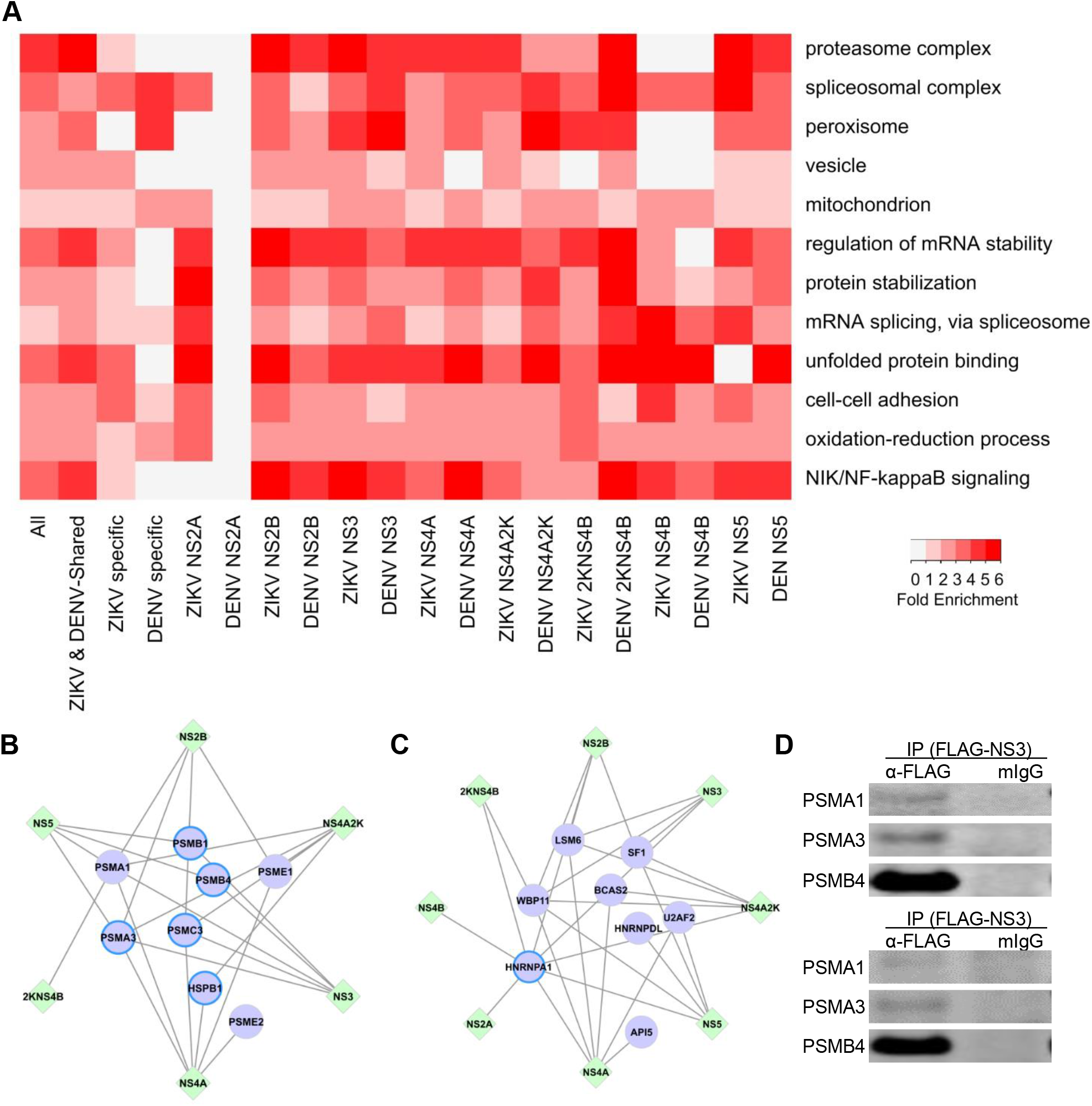
GO analyses of host proteins in the PPI networks. **A.** Enriched GO terms in the categories of Molecular Function, Biological Process, and Cellular Component are found in both shared and virus-specific PPI networks. The folds of enrichment are color-coded by *p* value. As examples, Interactions of six non-structural ZIKV proteins (NS2B, NS3, NS4A, NS4A+2K, 2K+NS4B, and NS5) with proteasome complex (**B**) and spliceosome complex (**C**) were shown respectively. Here, only the subunits capable of binding with ZIKV proteins were included in the figures. Circles with bright blue outlines indicate previously reported virus binding proteins. **D.** Co-IP of overexpressed FLAG tagged ZIKV-proteins and V5 tagged human proteasome subunits in 293FT cells. IP were performed with anti-FLAG mAb magnetic beads and eluted fractions were analyzed by Western blot using mouse anti-V5 antibody. Mouse IgG magnetic beads were used as a negative control to evaluate the non-specific binding on the beads. Inputs correspond to 2% of total lysate incubating with anti-FLAG mAb magnetic beads.

### Host cellular machineries involved in ZIKV-/DENV-human PPIs

To compare PPIs between ZIKV and DENV, we assembled a global PPI network involving 557 human, eight ZIKV, and eight DENV protein nodes (Fig.2D and Table S2). 147 and 42 host proteins were exclusively connected to either ZIKV or DENV proteins, respectively, suggesting that these virus-specific PPIs could contribute to ZIKV or DENV-specific infection outcomes or pathological effects. Interestingly, some human proteins exclusively interacted with a specific ZIKV protein, but not the homologous DENV protein. For example, PLEK connected only to ZIKV-NS2A, two human proteins, DDX49 and TTR, only to ZIKV-NS4B, and 75 proteins only to ZIKV-NS4A. Our recent study also confirmed interactions of ARPC3 and NUMBL to ZIKV-NS2A, but not to DENV-NS2A with co-IP method in HEK293 cells [27].

368 (66.1%) human proteins were connected to both ZIKV and DENV proteins in the PPI networks, supporting the notion that these two related viruses exploit similar cellular machineries. Statistical analysis showed a significant overlap between human proteins recognized among each viral homologous protein pairs (Figure 1C). For example, ZIKV-NS3 and DENV-NS3 proteins were found to interact with 238 and 240 human proteins, respectively, of which 187 were shared (78-79%; hypergeometric *p*-value = 3.7e-324). Conversely, ZIKV-NS3 and ZIKV-4A, two unrelated proteins, interacted with 238 and 401 human proteins, respectively, of which only 168 overlapped (42-71%). Similarly, only 127 proteins (53-68%) overlapped between 240 DENV-NS3 bound and 188 DENV-4A bound human proteins.

Gene Ontology (GO) analysis for human proteins that were targeted by each individual viral protein revealed several interesting features (**Figure 2**A, Table S3). First, host proteins connected to homologous ZIKV and DENV proteins were often enriched for the same GO terms, which is consistent with the result that a large number of shared host proteins interacted with homologous viral proteins. Second, host proteins targeted by different viral proteins were enriched for diverse biological processes and protein complexes. Third, many different viral proteins interacted with different components of the same enriched biological processes and protein complexes.

These observations raised the question of whether the conserved and virus-specific PPIs reflected different biological processes. Indeed, GO analysis of ZIKV/DENV conserved PPIs and ZIKV- or DENV-specific PPIs demonstrated distinct enrichments (Figure 2A). For instance, GO term of cell-cell adhesion was enriched mainly in human proteins specifically targeted by ZIKV proteins. On the other hand, GO terms of proteasome and NIK/NF-kappaB signaling were enriched in PPIs shared by ZIKV and DENV, suggesting that these virus-relevant biological processes may be important for flavivirus infections. For example, six of the non-structural ZIKV proteins (NS2B, NS3, NS4A, NS4A+2K, 2K+NS4B, and NS5) interact with eight components in the proteasome complex (Figure 2B). Similar phenomena were observed for the spliceosomal complex (Figure 2C). Furthermore, Co-IP was performed in HEK 293 cells to test the physical binding of proteasome subunit PSMA1, PSMA3 and PSMB4 to ZIKV-NS3 and NS5 respectively (Figure 2D). Consistent with our finding, a recent study reported that DENV-NS5 protein interfered with host mRNA splicing through direct binding to proteins in the spliceosome complex [33].

### RNAi screening identified critical host proteins for ZIKV replication

To validate whether host proteins enriched in the above biological processes and protein complexes were functionally involved in ZIKV infection, we carried out a siRNA knockdown assay similar to those used for other viruses [12, 34–37]. Specifically, 10,415 druggable target genes were individually knocked down and ZIKV NS1 protein level was measured using a high-throughput homogenous time-resolved fluorescence (HTRF) assay as a surrogate for viral load in ZIKV-infected HEK 293 cells. Among the 10,415 target genes, knockdown of 120 (1.2%) genes resulted in significantly reduced (> 30%) NS1 levels (Table S4). GO analysis revealed that proteasome, spliceosome, RNA polymerase, COPI vesicle coat and Eukaryotic 43S preinitiation complex were significantly enriched among those 120 genes with proteasome showing the lowest *P* value *(P* = 3.8e-25; **Figure 3**A).

**Figure 3.**
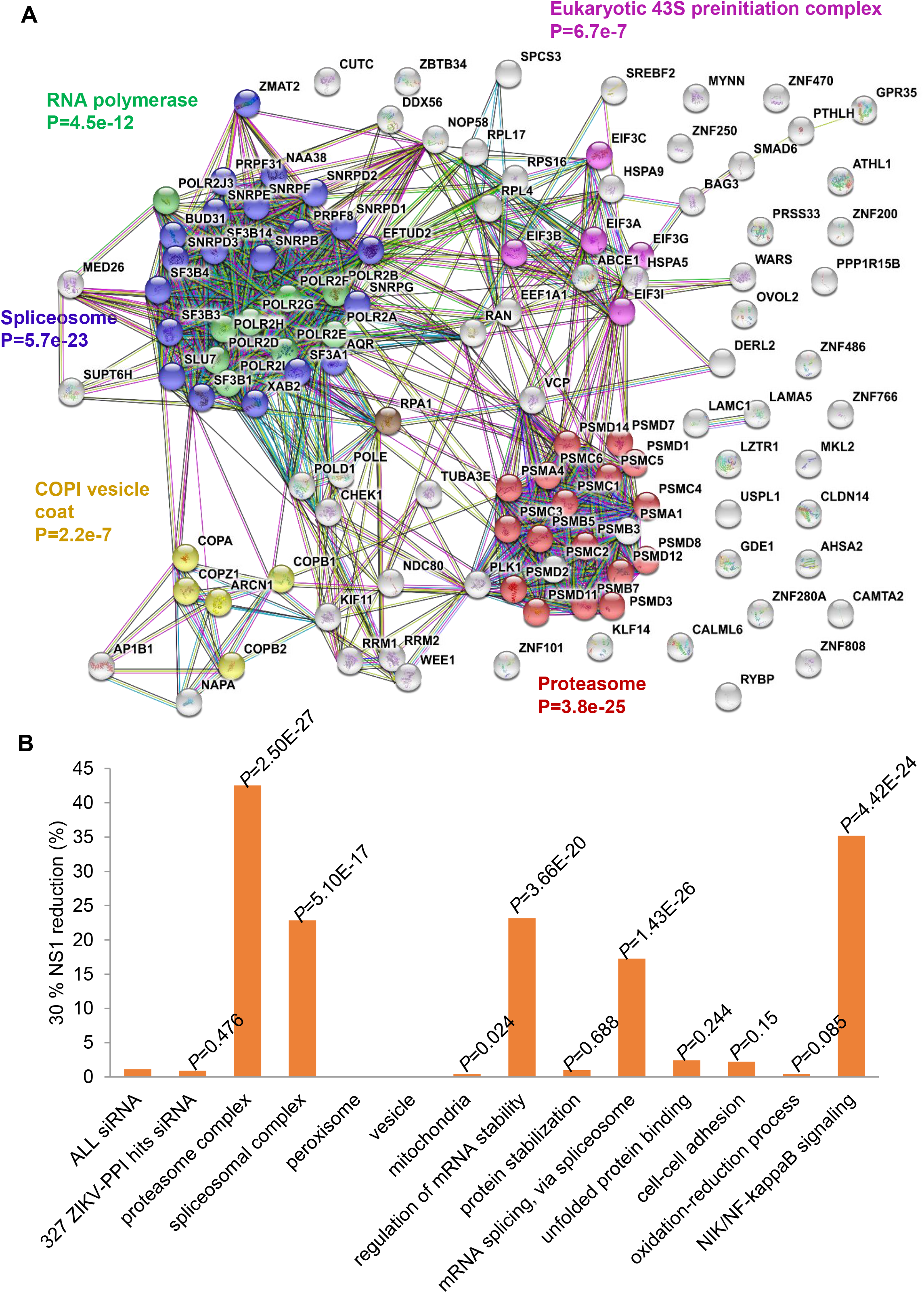
Identification of critical host proteins for ZIKV replication by RNAi screening. **A.** STRING analysis of genes that significantly affected ZIKV replication in RNAi screening revealed that proteasome, spliceosome, RNA polymerase, COPI vesicle coat and Eukaryotic 43S preinitiation complex were significantly enriched and proteasome showing the lowest *P* value of 3.8e-25 (FDR-adjusted). **B.** Shown are percentages of genes with over 30% reduction of NS1 levels by siRNAs among all genes in a specific category. The collection of siRNAs targets a total of 10,415 druggable genes (all siRNA group). Proteins produced by 327 genes interact with ZIKV proteins in the PPI dataset (ZIKV siRNA group). Note the high success rate (20 out of 47 members) in the category of “Proteasome Complex”.

Of the 10,415 target genes, protein products of 327 genes were found to interact with ZIKV proteins during our PPI analysis. Individual siRNA-knockdown of three (i.e., *PSMC3, PSMA1* and *OVOL2)* of them resulted in > 30% reduction of NS1 levels. Notably, a significant increase in the success rate of the knockdown assays was observed for those genes whose protein products were found in the enriched GO terms identified by the PPI analysis (Figure 3B). For example, individual knockdown of 20 of the 47 members in the proteasome complex showed >30% reduction of NS1 levels (42.5%; *P* = 2.5e-27; Figure 3B).

### High-throughput drug screening identified small molecule inhibitors

To further substantiate our results, we employed an independent chemical genetic approach to screen for and validate chemical compounds that target host proteins essential for viral replication and interfered with the ZIKV life cycle. A total of 6,016 compounds, including the Library of Pharmacologically Active Compounds (LOPAC, 1,280 compounds), the NIH Chemical Genomics Center (NCGC) pharmaceutical collection of 2,816 approved drugs, and 1,920 bioactive compounds [38], were screened for antiviral activity against ZIKV infection of HEK 293 cells. ZIKV infection was quantified by ZIKV-NS1 antibody-based Time-Resolved Fluorescence Resonance Energy Transfer (ZIKV NS1 TR-FRET) detection [39]. The ZIKV-NS1 TR-FRET assay measures the total amount of intra- and extracellular NS1 protein levels in infected cell culture, which was used as an indicator for ZIKV replication levels in cells (**Figure 4**A). Of the 6,016 compounds, 256 were identified as preliminary hits and selected for secondary validation by the NS1 TR-FRET assay and cytotoxicity evaluation in the same cells (Figure 4A). Viral inhibition was confirmed for 217 of the preliminary hits and 134 compounds exhibited greater than four-fold selectivity of ZIKV NS1 inhibition over compound cytotoxicity (Figure S3 and Table S5), which included all the 24 compounds previously reported [40].

**Figure 4.**
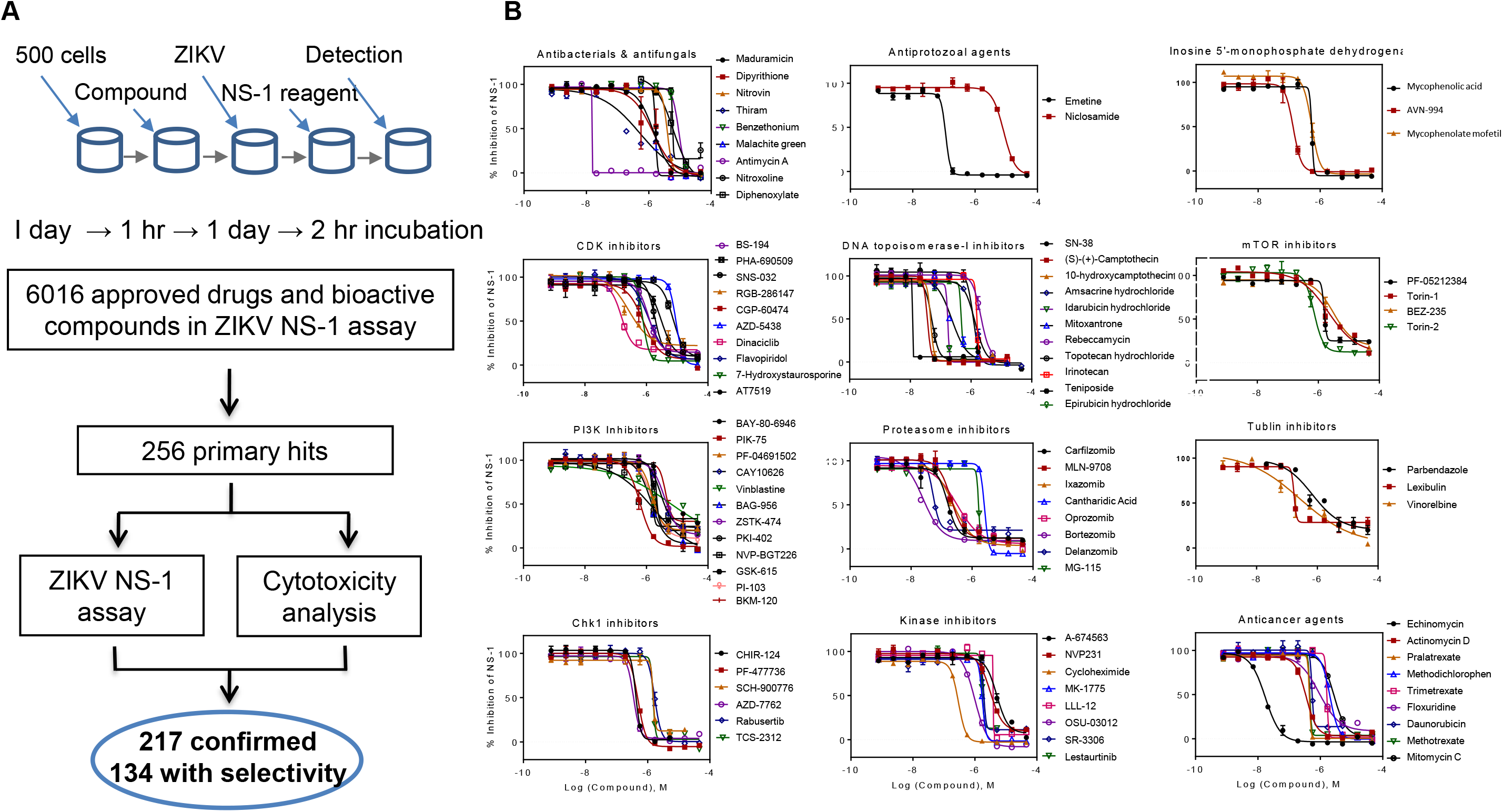
Identification of small molecule inhibitors against ZIKV replication. **A.** Flowchart of compound screening and confirmation with the NS1 assay. 6,016 drugs and bioactive compounds were added to precultured cells in 96-well plates, after infected with virus for one day and then ZIKV-NS1 TR-FRET assay were applied to measure the total amount of intra- and extracellular NS1 protein levels in the culture. Of the 6,016 compounds, 256 were identified as preliminary hits and selected for secondary validation by the NS1 TR-FRET assay and cytotoxicity evaluation with the same cells. 217 of the preliminary hits was confirmed and 134 compounds exhibited greater than four-fold selectivity of ZIKV NS1 inhibition over compound cytotoxicity. **B.** Summary of behaviors and IC_50_ values of 12 groups of potent compounds categorized based upon their reported mechanisms of action. Values represent mean + SD (n = 3 cultures). Curves represent best fits for calculating IC_50_.

Based on the reported mechanisms of action (https://tripod.nih.gov/npc/), ZIKV inhibition exhibited by 92 of 134 effective compounds was mainly mediated by 12 biological categories[38]: proteasome, antibacterials & antifungals, CDK inhibitors, PI3K inhibitors, Chk1 inhibitors, antiprotozoal agents, DNA topoisomerase-I inhibitors, kinase inhibitors, inosine 5’monophosphate dehydrogenase inhibitors, mTOR inhibitors, tubulin inhibitors, and anticancer agents (Figure 4B). To further confirm the anti-ZIKV activity of these compounds, the antiviral potency of effective compounds was determined using a ZIKV virus titer assay. Among these compounds, the antiviral activity of emetine was confirmed in the mouse models of ZIKV infection [41], which validated our compound screening approach. All results of the primary screen of the approved drug collection and hit confirmation were deposited into the open-access PubChem database under the ID: 1347053 (https://pubchem.ncbi.nlm.nih.gov/assay/assay.cgi?aid=1347053).

### Integrative analysis of omics data from PPIs, RNAi screen, and chemical genetics

We compared data deposited in Drugbank [42], Therapeutic Target Database (TTD) [43], and STITCH 5.0 [44] and identified 1,065 human proteins as targets of the 134 effective anti-ZIKV compounds from our screen. Of the 1,065 protein targets 45 were found to interact with ZIKV proteins in our PPI analysis (**Figure 5**A). STRING analysis revealed that the majority (80.0%) of these proteins were highly connected via functional associations, such as physical interactions, co-expression, tissue specificity, and functional similarity [45]. Indeed, 46 connections were found among 45 proteins, compared to only 18 expected connections (PPI enrichment *P* value = 1.3e-8). GO analysis of these proteins revealed significant enrichment in proteasome, vesicle and regulation of cell death (Figure 5A and Table S6).

**Figure 5.**
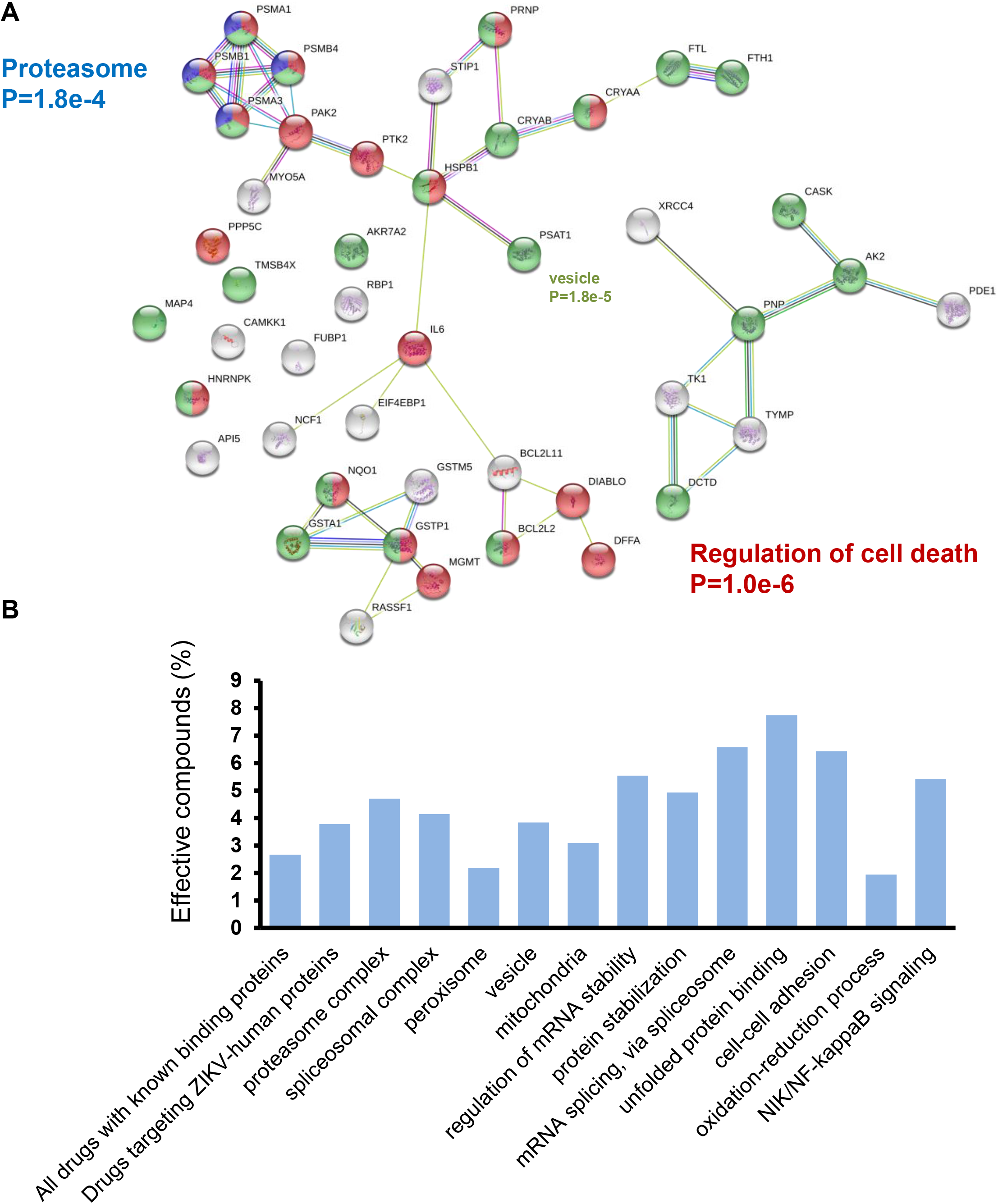
Integrative analysis of PPI and chemical genetics screen. **A.** 45 anti-ZIKV drug target human proteins were found to interact with ZIKV proteins in our PPI analysis. 80.0% (36/45) were highly connected via functional associations, such as physical interactions, co-expression, tissue specificity, and functional similarity. GO analysis of these proteins revealed significant enrichment in proteasome (FDR-adjusted *P* value of 1.8e-4), and regulation of cell death (FDR-adjusted *P* value of 1.0e-6). **B.** Functional association networks among the proteins that interact with viral proteins and are targeted by effective compounds.

Among the 6,016 tested compounds, there are 3,671 compounds with known targets. 98 of the 3,671 compounds (2.67%) showed selective inhibition against ZIKV infection. For the 766 drugs that are known to target proteins in our PPI analysis, 29 (3.79%) were effective, demonstrating a 1.42-fold enrichment (hypergeometric *p*-value = 0.02). Individual pathways and complexes also showed enrichment for identifying effective drugs, except for peroxisome and oxidation-reduction process (Figure 5B).

### Proteasome inhibitors suppress ZIKV and DENV replication

The integration of the three orthogonal datasets presented strong evidence that the same conserved cellular machineries play an important role in ZIKV and DENV replication. The proteasome complex stood out for several reasons. First, the PPI network analysis revealed that six ZIKV and six DENV proteins interacted with eight and seven proteasome subunits, respectively, most of which are part of the 20S core particle (**Figure 6**A-B). Second, individual knockdown of 20 proteasome genes resulted in substantially reduced ZIKV replication in the RNAi screen (Figure 3B). Third, the proteasome complex was the second most significantly enriched pathway targeted by the 134 effective compounds identified by the chemical genetics approach to inhibit ZIKV.

**Figure 6.**
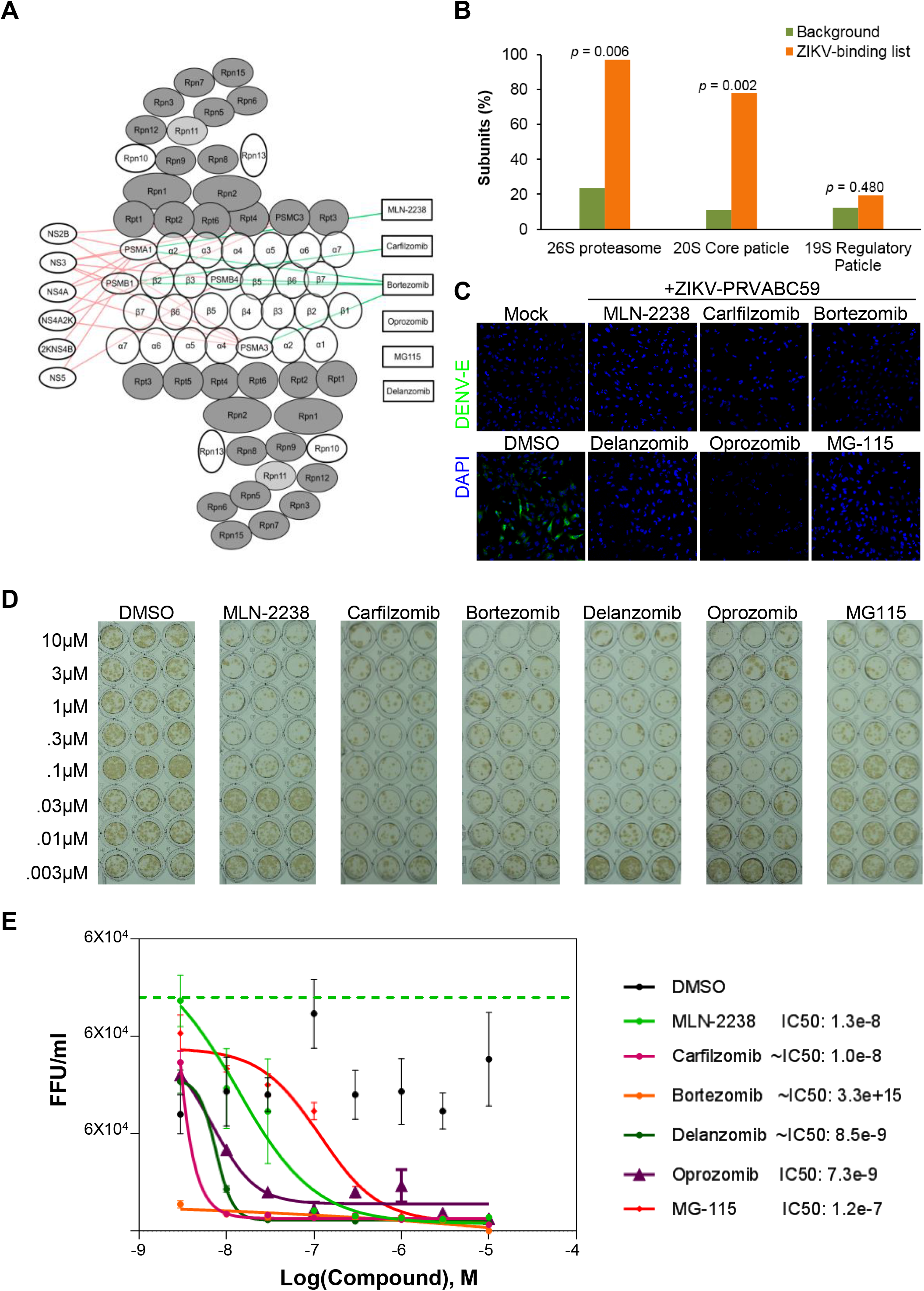
Inhibition of ZIKV expression and production by proteasome inhibitors. **A-B.** PPI network analysis of virus proteins and human proteasome subunits reveals that most of the interacting proteasome subunits are part of the 20S core particle. Percent of the binding subunits in 26S proteasome and its two sub-complexes, the 20S core particle and the 19S regulatory particle were presented. **C.** Inhibition of ZIKV expression in glioblastoma cells by a panel of proteasome inhibitors. The SNB-19 cells were infected by ZIKV PRVABC59 (MOI = 1) in the presence of 1 μm of each inhibitor and then incubated for 48 hrs before the cultures were analyzed for ZIKV envelope protein expression by immunostaining. Scale bar: 100 μm. **D-E.** Sample images (D) and quantification (E) of titer assay to assess the potency of the proteasome inhibitors against infectious ZIKV production in SNB-19 cells. All data were normalized to that for 0 μM for each compound. Dose dependent antiviral activity presented as fluorescent focus forming units (FFU/mL) and data represent mean + SD (n = 6). Curves represent best fits for calculating IC_50_ values (listed to the right).

To further validate our results, we selected six proteasome inhibitors (MLN-2238, Carfilzomib, Bortezomib, Delanzomib, Oprozomib, and MG-115) for further evaluation of their inhibitory activities on ZIKV and DENV in the human glioblastoma cell line, SNB-19. We used a recent clinical isolate of the Puerto Rico PRVABC59 ZIKV strain for this analysis. The cultures were infected with ZIKV or DENV at a multiplicity of infection (MOI) of 1 in the presence of these compounds at a concentration of 1 μM, with DMSO and niclosamide [39] serving as the negative and positive controls, respectively. All of the proteasome inhibitors tested suppressed both of ZIKV and DENV envelope expression in this assay as compared to the DMSO control (Figure 6C and Figure S4A-B).

Finally, we used a colorimetric focus-forming unit assay to determine the dose response and IC_50_ of these compounds on ZIKV production. Consistent with the intracellular antigen expression assay, all six proteasome inhibitors reduced infectious ZIKV production, with IC_50_ values for Carfilzomib, Bortezomib, Delanzomib, and Oprozomib in the nanomolar range (Figure 6D-E).

## Discussion

In this study we employed three high-throughput platforms to investigate host cellular machineries that are critical for ZIKV and DENV replication. First, HuProt arrays were used to screen for direct PPIs between each ZIKV/DENV protein and 20,240 human proteins. Next, a RNAi screen targeting 10,415 druggable genes was adapted to identify the critical human genes required for ZIKV replication. Last, a chemical genetics approach was employed to screen 6,016 bioactive compounds for their ability to inhibit ZIKV replication. We have confirmed the anti-ZIKV activities of 217 compounds with 134 of them having greater than 4-fold for the selectivity index which represents a comprehensive list of approved drugs and bioactive compounds with anti-ZIKV activity. Integration of the three independent omics datasets identified several host machineries, including the proteasome complex, the spliceosome complex, and regulation of mRNA stability. The integrated data, including PPI, RNAi screening, and compound screening in this study focused on ZIKV and DENV, provides useful resources for further studies to understand viral biology, disease pathogenesis and identify new drug targets. Moreover, the systematic screening illustrated by our approach can be readily implemented to study other virus-host interactions to uncover the nuances of disease pathogenesis and discover novel therapeutic strategies.

Our multi-omics datasets could have many applications. As an example, we recently took advantage of the PPI dataset to understand molecular mechanisms underlying the differential pathogenic impact on host cells induced by ZIKV and DENV [27]. Consistent with the clinical phenotype that ZIKV, but not DENV infection, could lead to microcephaly, our functional screen showed that expression of ZIKV-NS2A, but not the DENV-NS2A, leads to reduced proliferation and accelerated depletion of cortical neural stem cells in both embryonic mouse cortex in vivo and cultured human forebrain organoids. To understand how these two very similar proteins lead to different consequences in the same host cells, we mined the PPI dataset (Table S2) and found differential interactions of ZIKV-NS2A and DENV-NS2A with adherens junction proteins, which we validated in neural stem cells with endogenous proteins [27]. This critical information generated the hypothesis that the differential impact of ZIKV-NS2A and DENV-NS2A on adherens junctions may underlie their differential impact on neural stem cell properties, which we tested and confirmed in both in vivo embryonic mouse cortex and in vitro human brain organoid models [27]. Other viral proteins have also been implicated in the pathogenesis of virus infection; for example, ZIKV-NS4A and ZIKV-NS4B cooperatively suppressed the Akt-mTOR pathway to inhibit neurogenesis and induce autophagy in Human Fetal Neural Stem cell [46]. Additionally, we found targeting multiple components of the same protein complexes/signaling pathways seems to be a reoccurring event in pathogen-host interactions. Take the spliceosome complex as an example (Figure 2C) host proteins API5 and BCAS2 were found to interact with ZIKV proteins NS4A and NS4A2K, respectively. It is an intriguing point that the same process/complex can be targeted by a pathogen at different points. It is conceivable that such “multivalency” interactions could serve as an effective means to ensure the robust hijacking of the host cell machinery by a pathogen. For example, in one of our previous studies, we observed that four conserved viral protein kinases, encoded by four different herpesviruses, could all phosphorylate 14 components of the DNA damage response pathways, such as TIP60, RAD51, RPA1 and RPA2, using in vitro phosphorylation assays on human protein arrays [17]. In-depth in vivo studies confirmed that these phosphorylation events played an important role in promoting viral DNA replication in all four viruses. In another study, we observed that a secreted protein kinase ROP18, encoded by Toxoplasma gondii, could phosphorylate multiple components in the MAPK pathway [29]. A third example is the observation that KSHV-encoded LANA protein could bind to all three components of the NER damage recognition/verification complex XPA–RPA (i.e., XPA, RPA1 and RPA2) [47]. Therefore, our virus-host PPI database can be used to explore both conserved and unique pathogenic processes induced by ZIKV and DENV in different cellular contexts in the future.

In this study, we focused the investigation of our datasets on viral replication to identify critical cellular machineries as candidate drug targets [19]. Using high-throughput drug screening to reveal hijacked host machinery, we identified potential antiviral compounds with a higher genetic barrier for virus to develop drug-resistance. In addition, we could potentially use these host-targeting drugs as broad-acting antivirals for closely related viruses, such as DENV and ZIKV, because of their substantially overlapped PPI networks with the host. Integrative analysis of independently identified pathways and PPI networks presents a strong case for the proteasome as conserved, critical machinery for ZIKA and DENV replication. The proteasome complex is a part of the ubiquitin-proteasome pathway and regulates many fundamental cellular processes [48]. Emerging evidence implicates the proteasome as a critical player in viral pathogenesis by modulating the function of viral proteins to favor viral propagation and evade the host immune response [49–51]. Until now, there have been few FDA approved antiviral drugs targeting intracellular host proteins, due to the potential side effects [9]. Notably, Maraviroc, a CCR5 receptor antagonist, has been approved as an antiretroviral drug for the treatment of HIV infection, which could prevent viral entry by blocking binding of viral envelope gp120 to CCR5 [52]. Several proteasome inhibitor drugs tested in this study, including carfilzomib and bortezomib, have been approved by the FDA for the therapy of various cancers, such as breast cancer, multiple myeloma, and Hodgkin’s lymphoma [53–56]. Consequently, these drugs could potentially be repurposed to further evaluate their efficacy and tolerance in a clinical setting as novel therapies for ZIKV and DENV infection.

In summary, we discovered a multitude of cellular pathways and protein complexes related to ZIKV and DENV infection by integrating three high-throughput systems biology methods – ZIKA/DENV-human PPIs, a druggable genes screen, and high-throughput chemical genetics screening. We identified the human proteasome as a conserved critical machinery for ZIKV and DENV replication with functional confirmation by pharmacological proteasome inhibitors. We also found a comprehensive list of 134 selective ZIKV inhibitors that span over 12 cellular pathways and mechanisms. Our study provides a rich resource of multi-omics datasets for future investigation of viral pathogenesis and drug development and highlights a systematic biological approach to investigate virus-host interactions.

## Materials and methods

### Viral cDNA preparation

The African prototype ZIKV strain MR766 and DENV serotype 1 Hawaii strain were used to infect mosquito cells, as previously described [57]. Lysates of virus-infected mosquito cells were prepared, and one microgram of the total RNA was used to prepare cDNA by Superscript III (Catalog No. 18080044, Thermo Fisher Scientific, Waltham, MA) for PCR templates.

### Gateway cloning and protein expression

Gateway cloning and protein expression were performed using the method as our previous publication [58]. In short, primer sets with the *attB1* or *attB2* sequences at the 5’and 3’-ends (Table S1) were designed to amplify the full-length viral genes, which were then cloned into Gateway Entry vector pDONR221 using the Gateway recombination reaction. (Catalog No. 11789021, Thermo Fisher Scientific, Waltham, MA). Each initial cloning was examined by BsrGI (Catalog No. R0575S, New England Biolabs, Ipswich, MA) digestion and Sanger sequencing. Then, each insert viral gene was shuttled into the yeast expression vector pEGH-A to carry out the protein expression.

### Protein labeling

The quality and quantity of each ZIKV and DENV proteins were determined using SDS PAGE gel electrophoresis, followed by Coomassie stain. Proteins that passed this quality control test were then labeled directly with NHS-tethered Cy5 dye (Catalog No. GEPA15101, Sigma-Aldrich, St. Louis, MO) on the glutathione beads. After quenching the dye molecules, the labeled protein was eluted and the quality of these purified proteins was examined on SDS page gels.

### Identification of virus-binding host proteins on HuProt arrays

PPI assay on the Huprot array and signal extraction of each spot were performed using the same methods described previously [27]. In short, The signal intensity (*R_ij_*) of a given protein spot (*i,j*) was generated as foreground signal (F_*ij*_) minus the corresponding background signal (B_*ij*_). the averaged R_*ij*_ from duplicate spots was defined as the signal intensity of the protein probe (R_p_). For the replicate samples, the signal profiles were quantile normalized to a merged profile. Using a similar method as described in our previous studies [59], the Z-score of each binding assay with a virus protein was computed based on the distribution of R_p_,

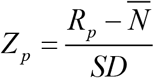

Where SD and N represent the standard deviation and mean of the noise distribution on the array, respectively. A stringent cutoff (Z ≥ 15) was used to call the positive hits in this study. The proteins determined as positives in all assays were removed for further analysis.

### Comparison to other datasets

The statistical significance of the overlap between our set of identified virus-binding human proteins and those deposited in VirusMint and Virhostome was calculated using hypergeometric test implemented in R [31, 32]. The number of background proteins was defined as the number of unique well-annotated human proteins detected in our HuProt Array (n = 13,816).

### Functional annotation

Database for Annotation, Visualization and Integrated Discovery (DAVID) was used to identify the enriched functional terms (molecular function, cellular component, biological process and KEGG pathway) for virus binding proteins [60]. Some enriched terms (P value < 0.05) were selected and represented in a heat map by the fold change.

### Protein-protein interaction network

Virus protein-human protein interactions identified in this study were input into Cytoscape to construct flavivirus-host PPI networks [61]. Human protein-protein interactions were extracted and drawn from STRING 10.0 [45]. The significance of functional terms and interaction numbers were also calculated and provided by STRING.

### Drug-target interaction

Drug targets were collected from three resources, Drugbank, Therapeutic Target Database (TTD) and STITCH 5.0 [42–44]. Drugbank and TTD include known targets of experimental drugs and FDA-approved drugs. STITCH combines chemical-protein interactions from experimental chemical screening, prediction, known database and text mining. For chemical-protein interactions in STITCH 5.0, only those with greater than 0.7 of high combined confidence score and with experimental or database scores were chosen for analysis. Those targets not identified as positive hits on HuProt arrays were removed for this study.

### Propagation of ZIKV

ZIKV stocks were generated in *Aedes albopictus* clone C6/36 cells as previously described [39]. PRVABC59-ZIKV strain was purchased from ATCC. MR766-ZIKV stock was purchased from Zeptomatrix. Briefly, a T-75 flask of C6/36 cells (90-95% confluency) was inoculated with 1 × 10^6^ ZIKV virions in low volume (3 mL) for 1 hr, rocking it every 15 mins. After 1 hr, 17 mL of media was added and C6/36 cells maintained at 28°C in 5% CO_2_ for 7 days. On day 7 and day 8 post-viral inoculation, supernatants were harvested, filtered, and stored at −80°C. ZIKV titer was determined by focus forming unit assay.

### Viral infections

For SNB-19 and Vero cell infections, cells were seeded into 12- or 96-well plates 1 day prior to viral infection. For SNB-19 cells, compounds were added 1 hr before addition of ZIKV at MOI = 0.5-1. SNB-19 cells were harvested at 24–48 hrs after infection for analysis by Western blot or immunofluorescence. For viral production assays, infected SNB-19 cell supernatant was harvested 24 hrs post infection, and analyzed by focus forming unit assay. ZIKV and DENV titers in cell supernatants were measured by focus forming units per ml (FFU/ml), as previously described [39].

### Co-immunoprecipitation and Western blot

FLAG tagged ZIKV-proteins (NS3 and NS5) and V5 tagged human proteasome subunits (PSMA1, PSMA3, and PSMB4) were overexpressed in 293FT cells. For Co-immunoprecipitation analysis, 293FT cells were incubated in the lysis buffer (Catalog No. #9803, Cell Signaling Technology, Danvers, MA) for half hour on ice. After sonication and centrifugation, the supernatants were subjected to immunoprecipitation with anti-FLAG mAb magnetic beads (Catalog No. M8823, Sigma-Aldrich, St. Louis, MO) at 4°C overnight. Then, the beads were washed for six times using lysis buffer and used to perform immunoblot assay with mouse anti-V5 antibody (Catalog No. R960-25, Thermo Fisher Scientific, Waltham, MA). Mouse IgG magnetic beads (Catalog No. #5873, Cell Signaling Technology, Danvers, MA) were used as a negative control to evaluate the non-specific binding on the beads. After incubating with Alex647 labeled Rabbit anti-mouse IgG secondary antibody (Catalog No. A-21239, Thermo Fisher Scientific, Waltham, MA) and washing, the membranes were visualized with Odyssey^®^ CLx Imaging System.

Additionally, for the western blot analysis of ZIKV and DENV proteins, SNB-19 cells were washed with PBS and directly lysed on ice in 1× Laemmli buffer. Lysates were then boiled and used for Western blot analysis. Membranes were probed with anti–ZIKV ENV (1:2,000; Catalog No. 1176-56, BioFront Technologies, Tallahassee, FL), anti-DENV NS3 (1:4,000; Catalog No. GTX133309, Genetex, Irvine, CA), or anti-GAPDH (1:20,000; Catalog No. sc-47724, Santa Cruz Biotechnology, Dallas, TX) and visualized with Odyssey^®^ CLx Imaging System after incubating with secondary antibodies.

### Immunocytochemistry

SNB-19 cells were seeded onto coverslips in 12-well plates 1 day prior to infection. At 24-hr post infection, cells were fixed with 4% paraformaldehyde for 15 mins at RT, followed by three 10-min washes in PBS at room temperature and permeabilized in PBT for 10 mins at room temperature (PBS with 0.1% Triton X-100). Cells were blocked for 1 hr at room temperature in PBTG, followed by incubation with anti–flavivirus group antigen 4G2 (1:1 000, Catalog No. ATCC^®^ VR-1852™, ATCC, Manassas, VA) at 4°C, washed three times with PBS, and incubated with goat anti-mouse-FITC (1:500, Catalog No. AP127F, Sigma-Aldrich, St. Louis, MO) for 1 hour at room temperature, followed by three subsequent 15 mins washes with PBS. Coverslips were mounted and nuclei stained using VECTASHIELD (Catalog No. H-1200, Vector Labs, Burlingame, CA).

### Compound screening using the TR-FRET NS1 assay

The primary compound screen was performed in 1536-well plates with the TR-FRET based NS1 assay as described previously [62]. Totally, there are 6,016 compounds were involved, including the Library of Pharmacologically Active Compounds (1,280 compounds, Catalog No. LO1280, Sigma-Aldrich, St. Louis, MO), NCGC pharmaceutical collection of 2,816 approved drugs, and 1,920 bioactive compounds [38].

For compound screening, HEK293 cells were seeded at 1,000 cells/well and incubated at 37°C with 5% CO2 for 16 hrs. Then, the compounds were transferred to cells in assay plates at 23 nl/well using a pintool workstation (Catalog No. NX-TR pintool station, Wako Automation, San Diego, CA) and incubated for 30 min. ZIKV (MOI = 1) was added to the assay plates at 2 μl/well followed by a 24-hr incubation. For detection of NS1 protein levels, 2.5 μl/well of TR-FRET NS1 reagent mixture was added and incubated overnight at 4°C. The plates were measured in the TR-FRET mode in an EnVision plate reader (Catalog No. 2105-0010, PerkinElmer, Waltham, MA). The experiment for hit compound confirmation was carried out in the same assay as the primary screen except all the compounds were diluted at a 1:3 ratio for 11 concentrations.

### Compound cytotoxicity assay

To eliminate the false positive compounds due to compound cytotoxicity, an ATP content assay [39] was used to measure cell viability after cells were treated with compounds in the absence of ZIKV-MR766 infection. Briefly, cells were plated in 1536-well white assay plates in the same way as described above. After a 24-hr incubation with compounds, 3.5 μl ATP content reagent mixture (Catalog No. 6016941, PerkinElmer, Waltham, MA) was added to each well in the assay plates and incubated for 30 mins at RT. Luminescence signals were determined in a ViewLux plate reader (Catalog No. ViewLux™ ultraHTS microplate imager, PerkinElmer, Waltham, MA). Compounds with cytotoxicity were eliminated from hit compound list as false positive compounds.

### siRNA screening

RNAi screening was conducted using the Ambion Silencer^®^ Select Human Druggable Genome siRNA Library Version 4 as described previously [63] and the HTRF assay for NS1 antigen was performed as described above. The HTRF signal was for each unique non-overlapping siRNA against the target genes was normalized to a negative control targeting siRNA. The values for each siRNA were divided by the median negative control values and multiplied by 100 to generate the negative normalized metric for each well/siRNA. The median value of negative controls in each plate were used for normalization, while the positive control was set to assess the assay performance and transfection efficiency.

## Data analysis and availability

The primary screening data and the curve fitting were analyzed as previously publication [64]. The concentration-response curves and IC_50_ values of compound confirmation data were analyzed using Prism software (https://www.graphpad.com/, GraphPad Software, Inc. San Diego, CA). Both of them were deposited into the PubChem database under the ID: 1347053 (https://pubchem.ncbi.nlm.nih.gov/assay/assay.cgi?aid=1347053).

## Author contributions

GS, EL, JP, and MX contributed equally to this work and were involved in experimental design, data collection and analyses. HR, YC, NW, SY, JK, CT, SM, CM, and KY contributed to additional experiments and data collection. KC, AS, WH, MX, RH, ML, HZ, HT, WZ, JQ, HS, and GM supervised the project. HT, WZ, JQ, HS, GM and HZ wrote the paper. ML, HT, WZ, JQ HS, GM and HZ were co-senior authors for the project.

## Acknowledgments

The authors would like to thank Paul Shinn and the compound management group at National Center for Advancing Translational Sciences (NCATS) for their professional support, and Jordan Schnoll for coordination. This work was supported by National Institutes of Health (U19AI131130, R01GM111514, R21AI131706, R35NS097370, and R37NS047344) and the Intramural Research Program of the NCATS/NIH.

## Supplementary material

**Supplementary Figure S1.**
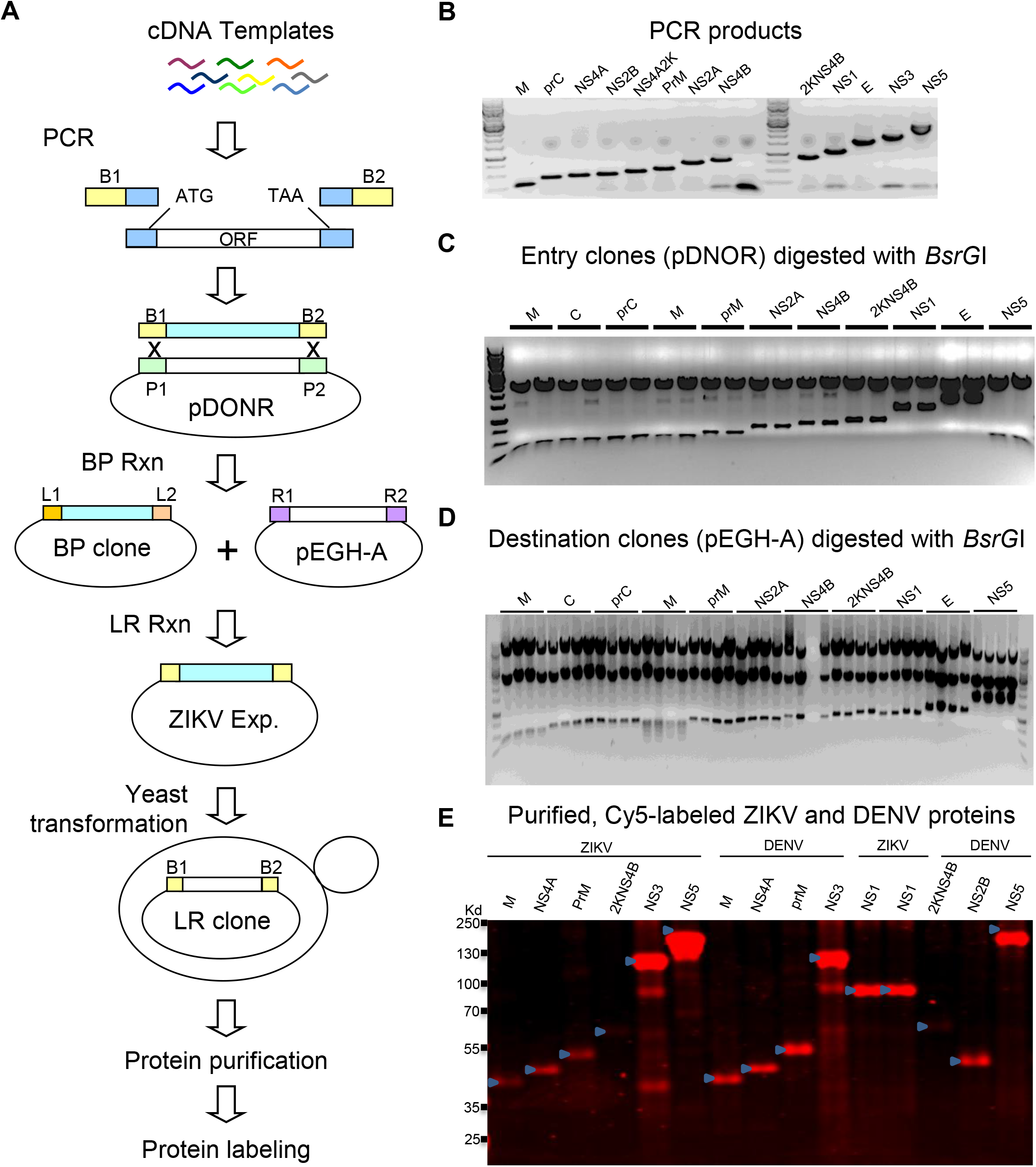
Preparation of fluorescent-labeled ZIKV and ENV proteins, **A.** Flow chart of fluorescent-labeled protein probes. **B.** Examples of successful PCR amplifications of ORFs using ZIKV cDNAs template. **C.** Examples of entry clones digested by BsrGI to release correct-size ORFs and then detected by 1% agrose gel. **D.** Examples of destination clones digested by BsrGI to release correct-size ORFs and then detected by 1% agrose gel. **E.** Examples of successful Cy5 labeled ZIKV and DENV proteins probes detected by SDS-PAGE gel. Blue arrows indicate each protein’s probe with correct molecular weight.

**Supplementary Figure S2.**
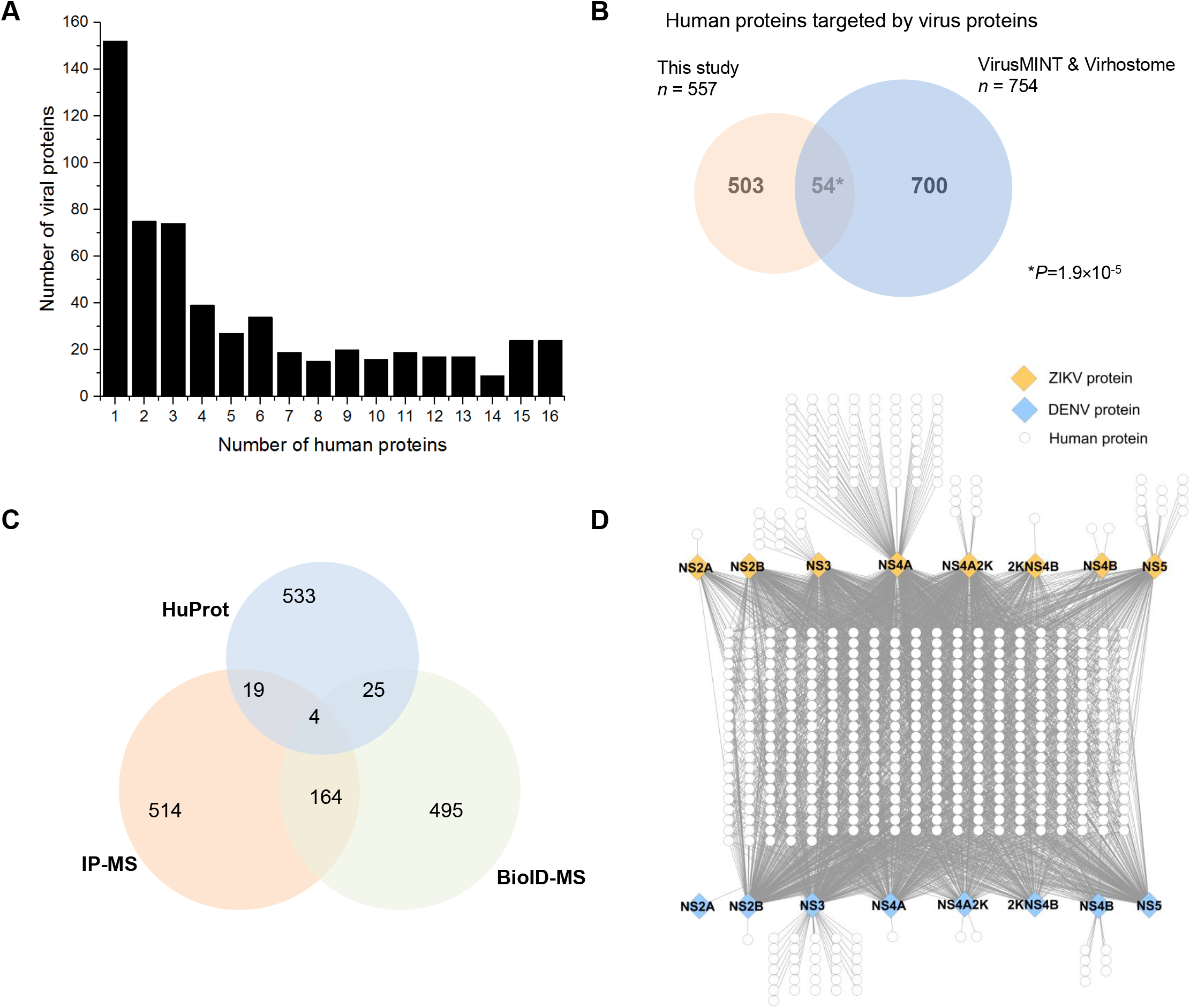
ZIKV- and DENV-host protein-protein binding interactomes. **A.** Interaction specificity of host proteins with viral proteins. Approximately 40% of human proteins identified by the PPI analysis only interacted with one or two viral proteins, while 24 human proteins interacted with all of the viral proteins tested and were removed from further analysis. **B.** Comparison of human proteins identified in this study with those known to be targeted by viruses in the VirusMINT and Virhostome databases. **C.** Comparison of human proteins identified in this study with a recently published data associated ZIKV-Human PPI based on MS method, which identified 701 vs. 688 human binding proteins by IP-MS and BioID respectively. **D.** The host proteins that interacted with both ZIKV and DENV proteins are shown in the middle of the PPI network, and the host proteins that interact specifically to either ZIKV and DENV proteins are placed on top or bottom of the network.

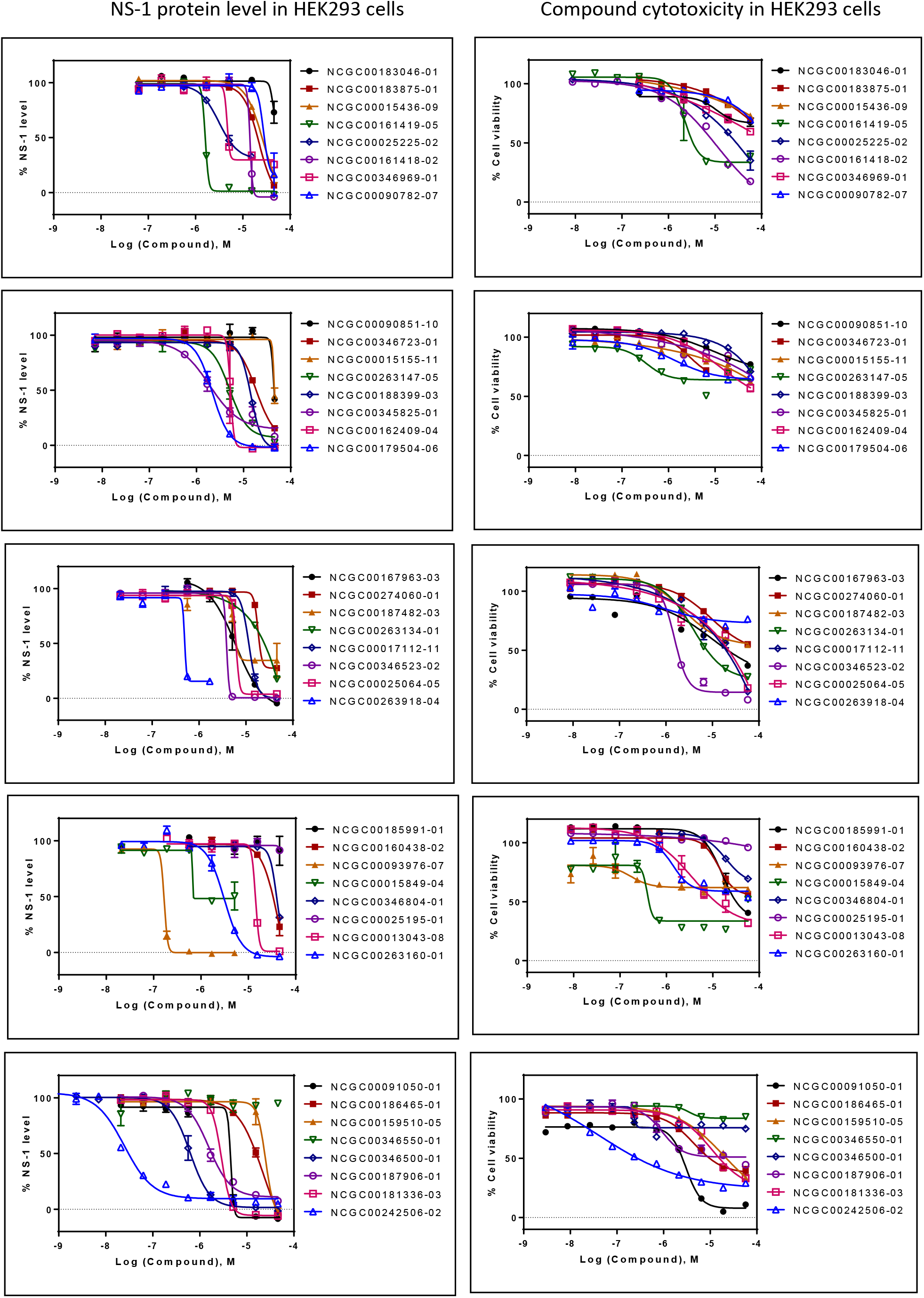
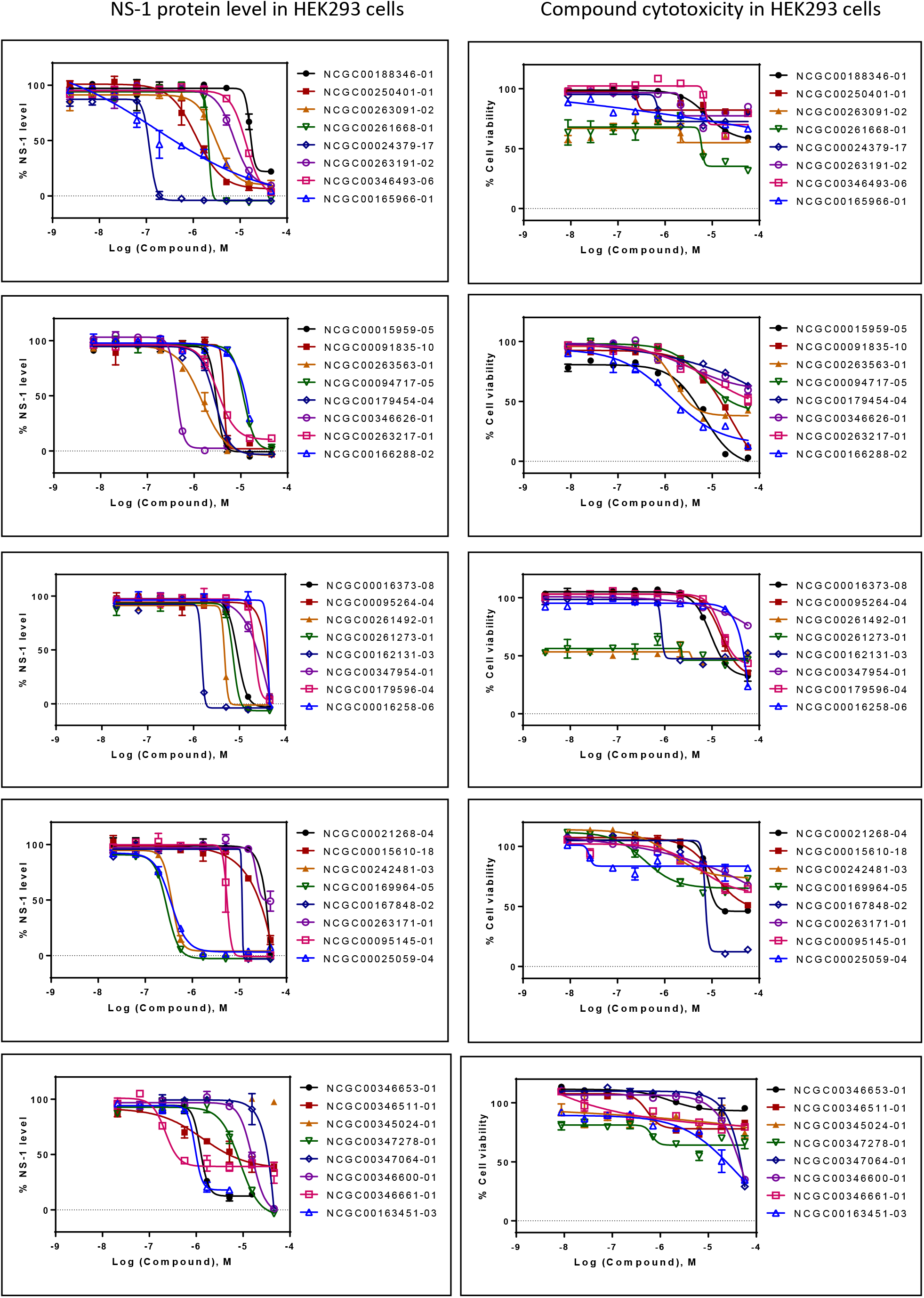
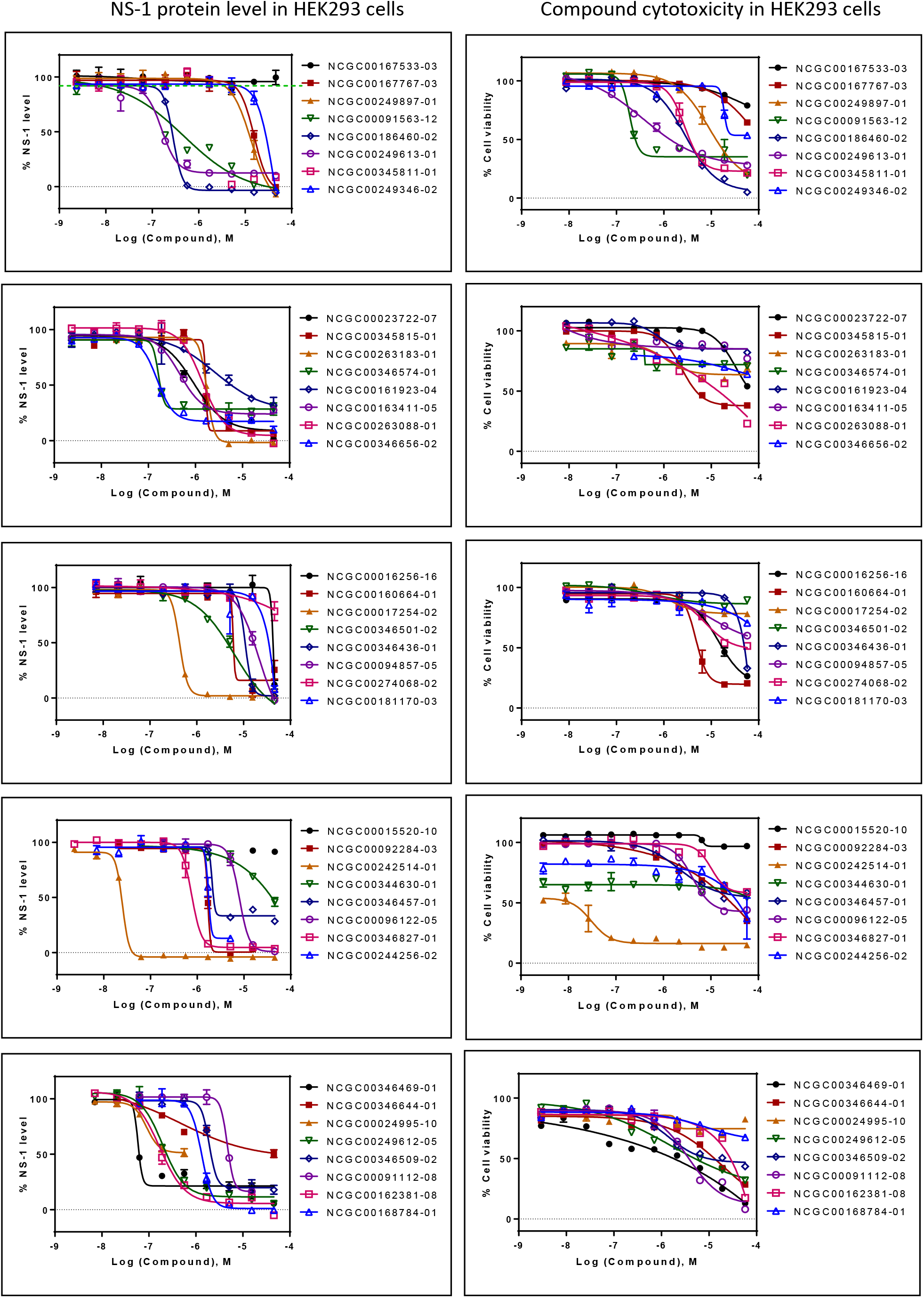
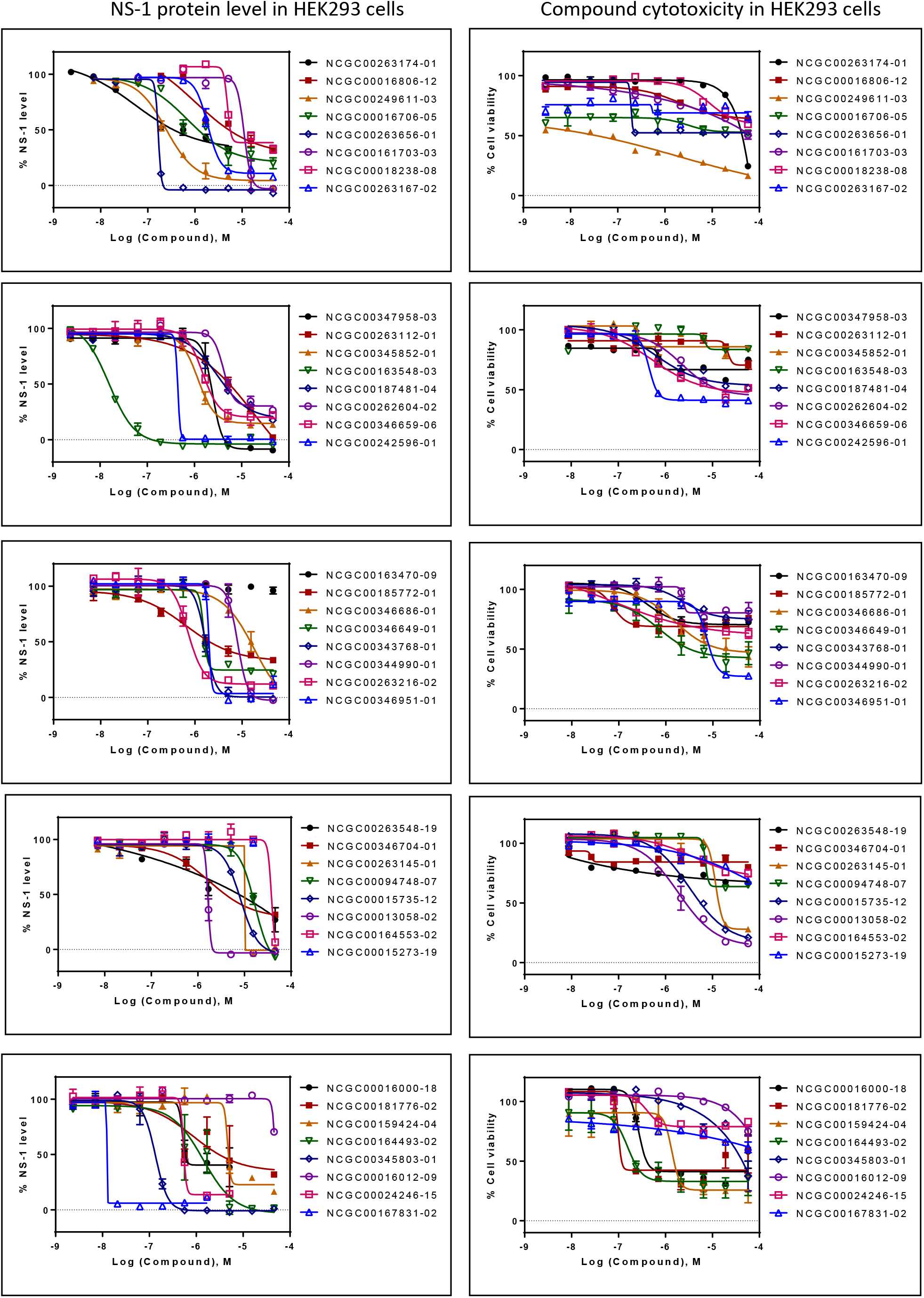
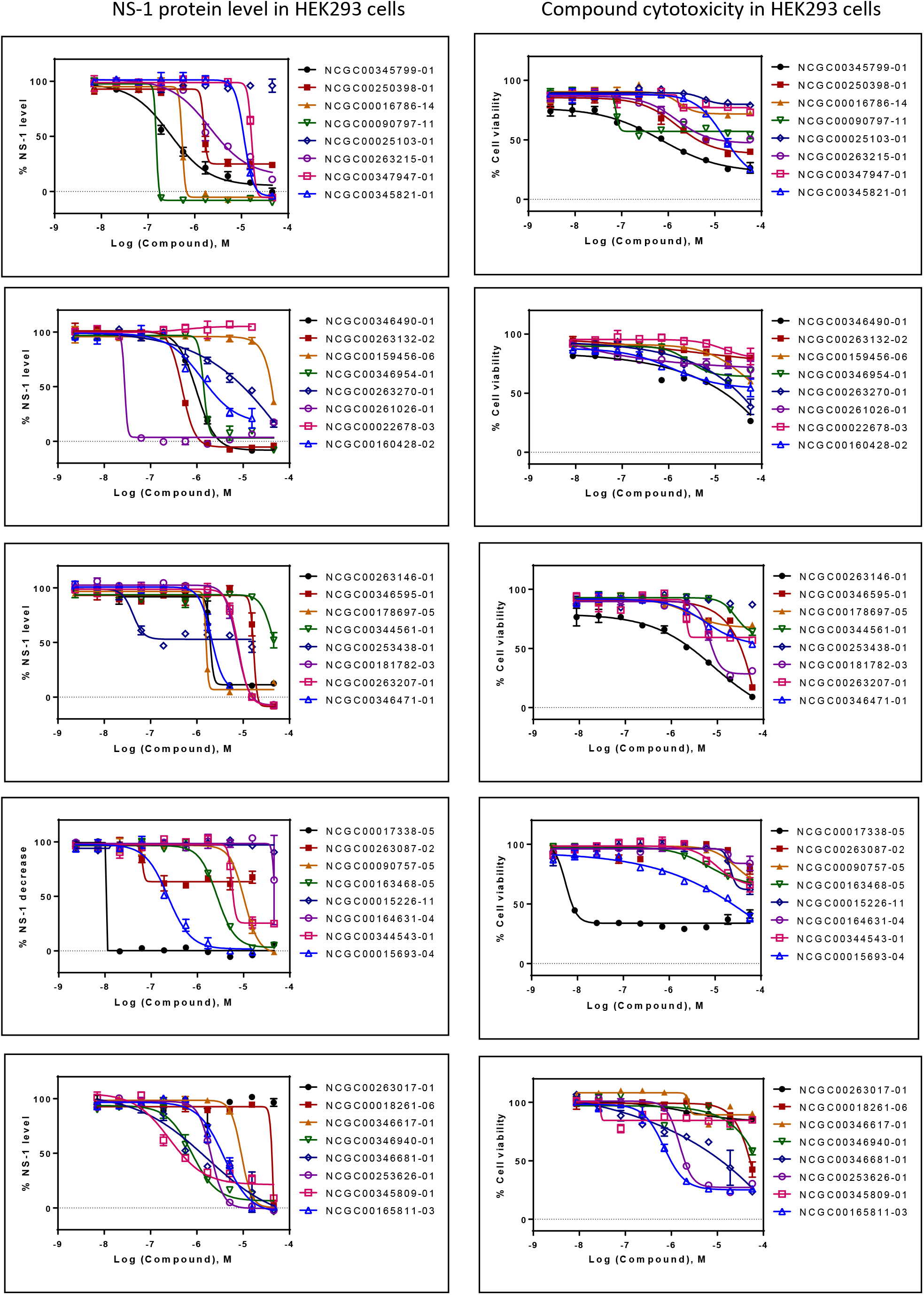
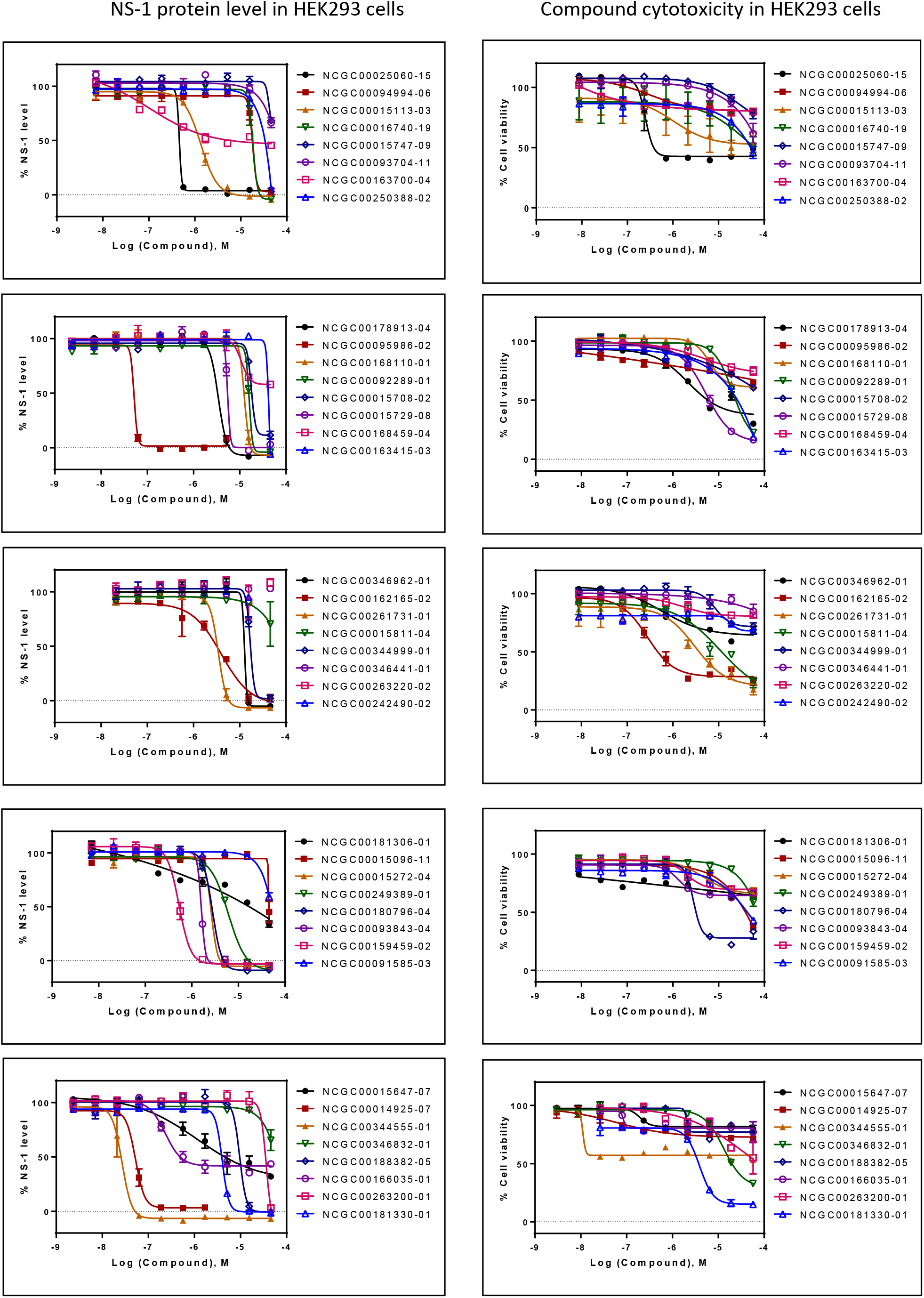
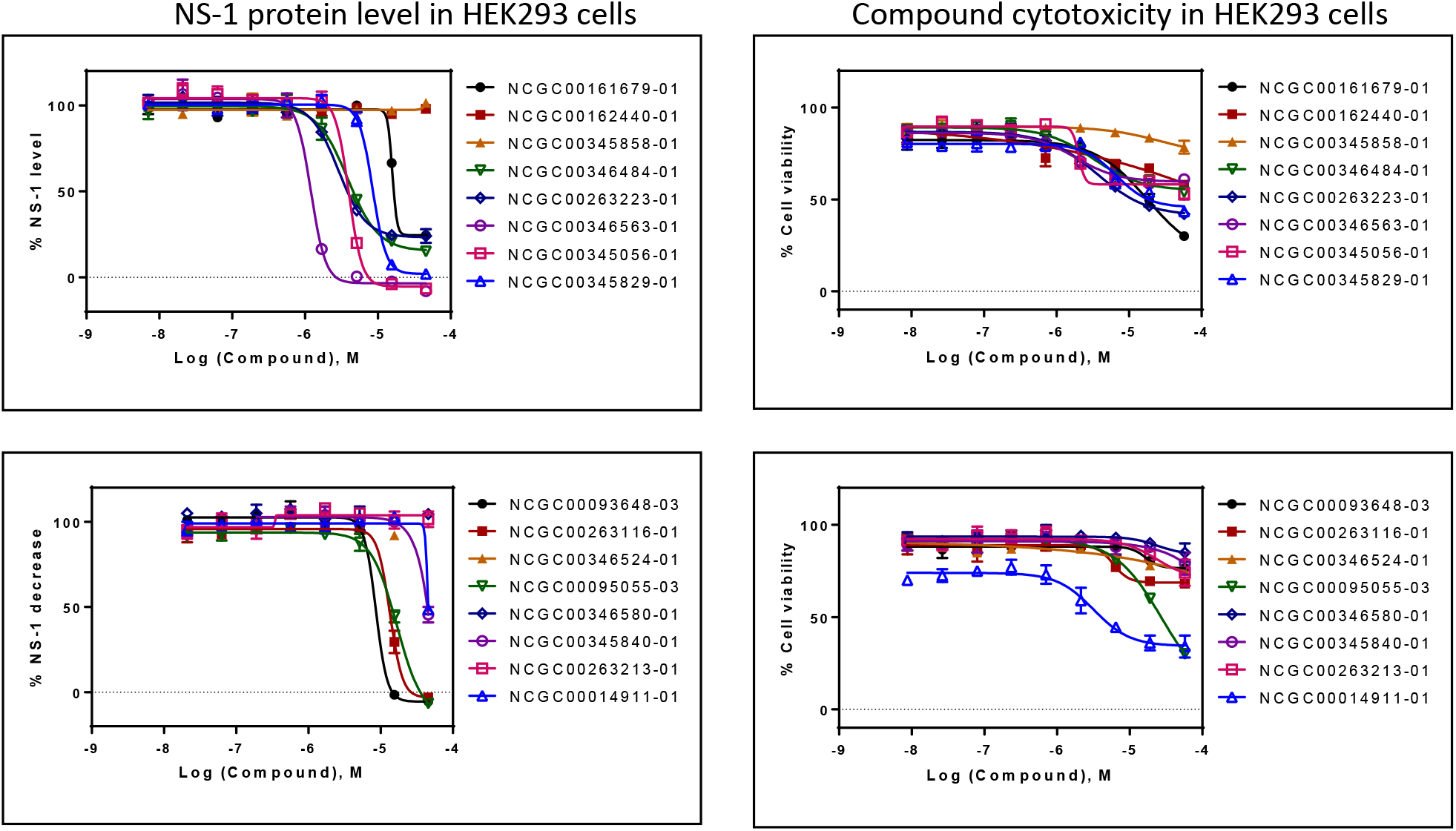

**Supplementary Figure S4.**
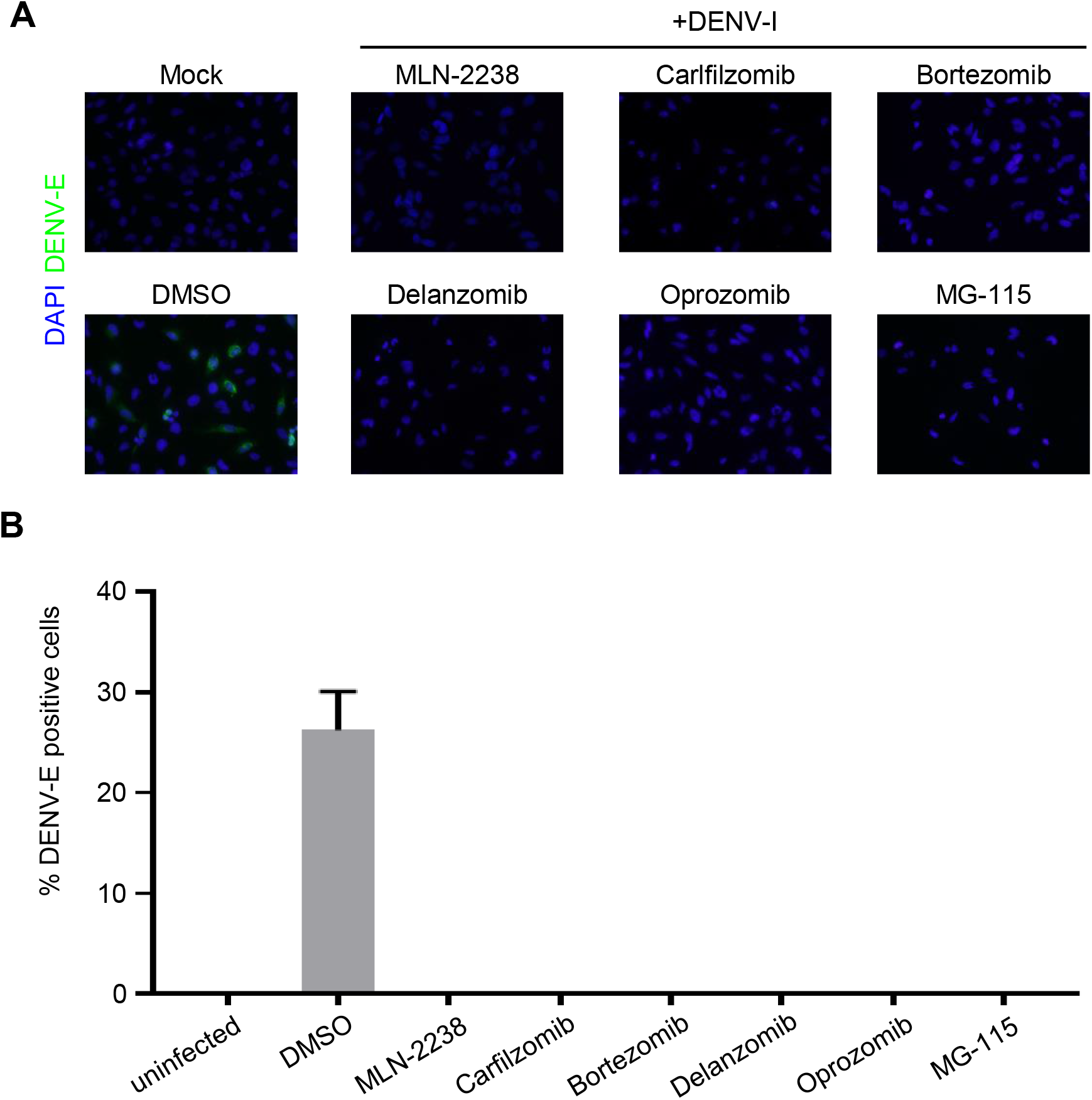
Suppression of DENV NS3 expression and viral production by proteasome inhibitors. **A.** The SNB-19 cells were infected by DENV (MOI = 1) in the presence of 1 μm of each inhibitor and then incubated for 48 hrs for immunocytochemistry of DENV envelope protein. **B.** Quantification of inhibition of DENV production by proteasome inhibitors as in (A). Values represent mean + SD (n = 3).

**Supplementary Table S1 PCR primers design for cloning of 13 ZIKV and 13 DENV full-length genes**

**Supplementary Table S2 Interactome of each individual ZIKV or DENV protein with human proteins identified on HuProt microarray**

**Supplementary Table S3 Entire list of enriched terms reported by DAVID**

**Supplementary Table S4 Summary of 120 genes with significant reduction of ZIKV in the siRNA knockdown assay**

**Supplementary Table S5 217 Active compounds inhibiting ZIKV NS-1 production in vitro**

**Supplementary Table S6 GO enrichment analysis carried out with STRING**

